# *Lactiplantibacillus plantarum K8* lysates regulate hypoxia-induced gene expression

**DOI:** 10.1101/2023.12.08.570897

**Authors:** Jaehyeon Jeong, Byeong-Hee Kang, Sangmin Ju, Na Yeon Park, Deukyeong Kim, Ngoc Thi Bao Dinh, Jeongho Lee, Chang Yun Rhee, Dong-Hyung Cho, Hangeun Kim, Dae Kyun Chung, Heeyoun Bunch

## Abstract

Hypoxic responses have been implicated in critical pathologies, including inflammation, immunity, and tumorigenesis. Recently, efforts to identify effective natural remedies and health supplements are increasing. Previous studies have reported that the cell lysates and the cell wall-bound lipoteichoic acids of *Lactiplantibacillus plantarum K8* (K8) exert anti-inflammatory and immunomodulative effects. However, the effect of K8 on cellular hypoxic responses remains unknown. In this study, we found that K8 lysates had a potent suppressive effect on gene expression under hypoxia. K8 lysates markedly downregulated hypoxia-induced HIF1α accumulation in the human bone marrow and lung cancer cell lines, SH-SY5Y and H460. Consequently, the transcription of known HIF1α target genes, such as *p21*, *GLUT1*, and *ALDOC*, was notably suppressed in the K8 lysate supplement and purified lipoteichoic acids of K8, upon hypoxic induction. In addition, K8 lysates decreased the expression of PHD2 and VHL proteins, which are responsible for HIF1α destabilization under normoxic conditions. Overall, our results suggest that K8 lysates desensitize the cells to hypoxic stresses and suppress HIF1α-mediated hypoxic gene activation.

## INTRODUCTION

Gene regulation is a fundamental and essential process through which living organisms develop, grow, maintain, and perish. Cells that constitute a living organism respond to the environmental and organismal signals and ensure proper gene expression based on the received signals. Hypoxia is an urgent condition that triggers rapid gene regulation in cells to manage low-oxygen stress^1^. DNA microarray and transcriptome profiling analyses have revealed that several hundred genes are differentially expressed upon exposure to hypoxic conditions^2,3^. These hypoxia-responsive genes are involved in diverse pathways including energy/glucose metabolisms, cell proliferation, and apoptosis^4–6^. Hypoxic stress is not only caused by an oxygen-deficient environment such as a high-altitude setting or anemia, but is also accompanied by certain physiological microenvironments, including tumors and inflammation^7^. For instance, hypoxic response is important for tumor progression and proliferation, and dozens of anticancer drugs that suppress hypoxic gene activation have been developed and preclinically administered^8,9^.

Hypoxia-inducible factor 1 (HIF1) consists of heterodimeric subunits with isoforms^6,9^. HIF1α is the major subunit and its expression is regulated by oxygen levels in the cell. Under normoxia, HIF1α in the cytoplasm is proline-hydroxylated and ubiquitinated for proteasomal degradation by prolyl hydroxylase (PHD) and von Hippel Lindau (VHL) proteins, respectively^10–12^. On the other hand, under hypoxic conditions, HIF1α is stabilized, translocated into the nucleus, and dimerized with constitutively expressed HIF1β^12^. The HIF heterodimer binds to the hypoxia response elements on its target genes to activate them. Therefore, the regulation of the HIF1α protein levels is critical to determine whether the cell suppresses or triggers hypoxic responses^13^. Notably, the over-expression of PHD proteins inhibits HIF1α accumulation, whereas HIF1α also reportedly induces PHD gene expression^14–16^, suggesting a possible feedback inhibition between the two enzymes. The VHL protein is known as a tumor suppressor and E3 ligase that ubiquitinates HIF1α for degradation^17,18^.

*Lactiplantibacillus plantarum K8* (K8) is a Lactobacillus strain isolated from a spicy, fermented cabbage dish called kimchi^19^. Several research groups have identified interesting health benefits of K8. K8 lysates (K8-L) or K8 lipoteichoic acid (LTA; K8-A) associated with the cell wall can modulate immunity and suppress inflammation and obesity^20–23^. The lipoprotein and LTA components of K8 appear to interact with toll-like receptor 2 (TLR2) in host cells to suppress inflammatory responses^24,25^. K8-L and K8-A appear to regulate the gene expression of the critical genes in these pathways^21,23,26^. For example, K8-L reportedly inhibits the expression of PPARγ and C/EBPα, which are important for adipogenesis, and the pre-inflammatory factors, interleukin (IL)-1β and -6 and TNF-α at the transcriptional level^21,25,27^. K8-A supplement effectively suppresses *Shigella flexneri* peptidoglycans-stimulated cytokine expression by dampening the activation of MAPK pathway including ERK and JNK in human monocyte-like cell line, THP-1^20^. It is not surprising that environmental substances, such as K8-L or K8-A, when treated on the cells or injected into the extracellular matrix, are capable of switching gene expression directly or indirectly through signal-transduction. Accumulating evidence has shown transcriptomic regulation by environmental and natural chemicals^28–31^. Understanding their physiological effects and gene regulatory mechanisms has been of interest because the applications of these natural chemicals, which are likely to have fewer adverse health effects, might replace those of synthetic chemicals.

Although K8-L and K8-A have been reported to modulate inflammation and immunity^21,25,32,33^, their effects on hypoxia, which is closely related to the inflammatory and immune responses, remain unknown. In addition, recent studies indicate that the chemicals released from gut bacteria can affect brain function and neurological health^34^ and lung function and infection^35^. We hypothesized that K8 cellular compounds might regulate cellular hypoxic responses through controlling HIF1α activation, with relatively low toxicity, and attempted to understand the effects of heat-treated K8-L (hK8-L) and K8-A on hypoxic gene expression in the human bone marrow and lung cancer cell lines, SH-SY5Y and H460, respectively. We hypothesized that HIF1α stabilization under hypoxic stress might be compromised by K8-L and if so, the expression of HIF1α target genes would be interfered to dampen hypoxic responses. We utilized cytotoxicity and molecular cell-based analyses to address these hypotheses. Our data suggest that hK8-L and K8-A are rarely cytotoxic and have potent suppressive activities against the stability of HIF1α and the activation of representative HIF1α-target genes under hypoxic stresses.

## MATERIALS & METHODS

### K8 sample preparation

*L. plantarum K8* (KCTC10887BP) was cultured in 1 L of MRS broth (BD Bioscience, USA) at 37 °C overnight and then cells were harvested by centrifugation at 8000× g for 8 min. Bacterial cells were washed with deionized water (DIW) and re-suspended in DIW and disrupted by a microfluidizer five times at 27,000 psi. Disrupted *L. plantarum K8* was freeze-dried to make *L. plantarum K8 lysates*. They were resuspended in PBS and the mixture was boiled at 100 °C for 20 min to inactivate live bacteria and proteins. Inactivation of live bacteria was confirmed by plating on MRS plates after heat treatment and no colonies were formed. PBS and PBS-soluble substances were removed by centrifugation. The PBS-insoluble pellet was dissolved in DMSO and designated hK8-L. The concentration of hK8-L was calculated as the weight of wet pellet divided by the volume of DMSO. Furthermore, hK8-L was centrifuged to remove the insoluble substances and debris, and the supernatant was filtered using 0.2 μm syringe filter to generate hK8-FL. Highly purified LTA was isolated from *L. plantarum K8* (KCTC10887BP) by n-butanol extraction, as previously described^36^. Briefly, L. plantarum K8 was cultured in 8L MRS broth for 16 h at 37 °C. The cells were harvested by cengtrifugation at 8000x g for 8 min, suspended in 0.1 M sodium citrate buffer (pH 4.7), and disintegrated by ltrasonication. The disrupted bacterial cells were then mixed with an equal volume of n-butanol by stirring them for 30 min at RT. After centrifugation at 13,000 ×g for 20 min, the aqueous phase was evaporated, dialyzed against pyrogen free water, and equilibrated with 0.1 M sodium acetate buffer containing 15% n-propanol (pH 4.7). The LTA was purified by hydrophobic interaction chromatography on an octyl-Sepharose CL-4B (Sigma) column (2.5 cm by 20 cm). The column was eluted with a stepwise n-propanol gradient (100 mL of 20% n-propanol, 200 ml of 35% n-propanol, and 100 mL of 45% n-propanol). Then, the column fractions containing LTA were pooled after an inorganic phosphate assay, and the pool was dialyzed against water. The purity of the purified LTA was determined by measuring the protein and endotoxin contents through the conventional silver staining after polyacrylamide gel electrophoresis and being through the Limulus amebocyte lysate (LAL) assay (pLTA<0.031 EU/ml) (BioWhittaker, U.S.A.). Nucleic acid contamination was assessed by measuring UV absorption at 260 and 280 nm.

### Cell culture and conditions

SH-SY5Y and H460 cells (American Type Culture Collection, USA) were grown in high glucose-DMEM (Cytiva, USA) and RPMI-1640 media (Gibco, USA), respectively, supplemented with 10% fetal bovine serum (FBS, Gibco, USA) and 1% penicillin/streptomycin (P/S, Thermo Fisher, USA) solution at 5% CO_2_ incubator at 37°C. The cells were grown to 70–80% confluence in a 10 cm dish before splitting into a 6 or 96 well plate. K8 lysates (DMSO, control) were applied according to indicated concentrations for 24 h. CoCl_2_ (Cat. 60818, Sigma, USA) or deferoxamine methanesulfonate (DFO; D9533, Sigma, USA) salt, was treated to indicated final concentrations (H_2_O, control) in the halfway, 12 h (CoCl_2_) or 24 h (DFO) after K8 supplement, for 12 h before collecting the cells for the assays.

### Cytotoxicity test

SH-SY5Y and H460 cells were grown to approximately 50–60% confluence in a 96-well plate in complete media. The media were exchanged with the fresh one including K8 lysates in DMSO, 1% of the total media volume. The K8 lysates-treated cells were incubated for 12 h before 100–200 μM CoCl_2_ in H_2_O or an equal amount of H_2_O, 1% of the total media volume. After an additional f12 h incubation, 10% of water-soluble tetrazolium salt (WST, DoGenBio, South Korea) was added to each well following the manufacturer’s instruction. The reaction was developed for 45 min to 1 h and the absorbance was measured at 450 nm using a microplate reader (Tecan, Switzerland).

### Immunoblotting

SH-SY5Y cells were grown to approximately 70-80% confluence in 6-well plates. The media were exchanged with the fresh one including K8 lysates in DMSO, to 1% total media volume. The K8- or DMSO-treated cells were incubated for 12 h before CoCl_2_ supplement to the designated concentrations. After an additional 12 h incubation, the cells were washed with cold PBS and collected with RIPA buffer (Cell Signaling Technology, USA). The protein concentrations of cell lysates were measured using the Bradford assay (Bio-Rad, USA). From the measured protein concentrations, equal amounts of proteins were loaded and ran on a SDS-polyacrylamide gel. The separated proteins were transferred to the nitrocellulose membrane (GE Healthcare, USA). Primary antibodies were used for probing HIF1α (#14179, Cell Signaling Technology, USA), p21 (sc-6246, SantaCruz Technology, USA), PHD2 (sc-271835, SantaCruz Technology, USA), β-ACTIN (MA5-15739, Invitrogen, USA), and α-tubulin (sc-8035, SantaCruz Technology, USA). Each antibody was diluted in a 5% skim milk (MBcell, South Korea or Bio-Rad, USA) solution or TBST solution [20 mM Tris-HCl, 137 mM NaCl, 0.1% (v/v) Tween-20, 5% (w/v) BSA, 0.025% (v/v) sodium azide and pH 7.6]. After the primary antibody incubation, the membranes were rinsed twice and washed four times, each for 10 min, with PBST solution [137 mM NaCl, 2.7 mM KCl, 4.3 mM Na_2_HPO_4_, 1.5 mM KH_2_PO_4,_ 0.1% (v/v) Tween-20 and pH 7.4]. The secondary antibodies, HRP-conjugated rabbit (#7074, Cell Signaling Technology) or mouse (#7076, Cell Signaling Technology), were diluted in 5% skim milk and incubated with the membranes. After incubation, the membranes were rinsed twice and washed with PBST solution four times, each for 10 min. Western Blotting Luminol Reagent (sc-2048, SantaCruz Technology, USA) or SuperSignal West Atto Ultimate Sensitivity Substrate (A38554, Thermo Fisher, USA) was used to develop signals. When indicated, immunoblotting images were quantified using Image J (NIH, USA).

### Chromatin immunoprecipitation (ChIP)

ChIP was performed as described in our previous studies ^37,38^. Briefly, the cell lysis buffer including 5 mM PIPES (pH 8.0), 85 mM KCl, and 0.5% NP-40 was used to resuspend the crosslinked cells. The cells were resuspended with nuclei lysis buffer including 50 mM Tris-HCl (pH 8.0), 10 mM EDTA, and 1% SDS immediately before sonication. Sonication was performed on ice at 22% amplitude for 30 s with 2 min intervals between pulses (Vibra-Cell Processor VCX130, Sonics, USA). The number of pulses was optimized to produce DNA segments ranging between 100 and 1000 bp on a DNA gel. Protease inhibitors were freshly added to the cell and nuclei lysis buffers including 1 mM benzamidine (Sigma, USA), 0.25 mM PMSF (Sigma, USA), aprotinin (1:1000, A6279, Sigma, USA), 1 mM Na-metabisulfite (Sigma, USA), and 1 mM DTT. The ChIP-grade magnetic beads coated with protein G were purchased from Cell Signaling Technology (USA). The antibodies used in immunoprecipitation were Pol II (#39097, Thermo Fisher, USA) and HIF1α (#14179, Cell Signaling Technology, USA). After IP and reverse cross-linking, DNA was purified through Qiagen PCR purification kit (Qiagen, Germany). The ChIP products were analyzed by real-time PCR using SYBR Green Realtime PCR Master Mix (Applied Biosystems, USA) under thermal cycling as 1 min at 95 ℃ followed by 45 cycles of 15 s at 95℃, 15 s at 55℃, and 1 min at 72℃. The results were analyzed as percent inputs. The primer sets amplifying the transcription start site and the gene body of the HIF1α target genes are listed in Table S1.

### Quantitative real-time PCR (qRT-PCR)

cDNAs were constructed from 600 ng of the collected RNAs by reverse transcription using ReverTra Ace qPCR RT Master Mix (Toyobo, Japan). Equal amounts of resultant cDNAs were analyzed through Real-time qPCR using Applied Biosystems PowerUp SYBR Green Master Mix and according to the manufacturer’s instruction (Applied Biosystems, USA). StepOnePlus Real-Time PCR System was used (Thermo Fisher Scientific, USA). The thermal cycle used was 10 min for pre-denaturation, followed by 45 cycles of 95°C for 15 s, 55°C for 15 s, and 72°C for 45 s. The primers used for the experiments were purchased from Integrated DNA technology (IDT, USA) and were summarized in **Supplementary Table 1.**

### Fluorescence-activated cell sorting (FACS)

Flow cytometric analysis was performed to determine the cell cycle status. The cells were harvested by trypsinization and fixed with ice-cold 70% ethanol for 15 min at –20°C. The cells were washed with PBS twice and suspended in 1 ml of the cold propidium iodide solution. Then the cells were incubated at room temperature for 15 min and analyzed by a flow cytometer (Attune NxT Flow Cytometry, Thermo Fisher).

### Statistical analysis

Standard deviation was calculated and used to generate error bars. The Student’s t-test was used to determine statistical significance (*P* < 0.05, one-sided). Graphs were generated using the Prism 8 software (GraphPad, Inc.).

## RESULTS

### hK8-L suppresses HIF1α stabilization under hypoxia

K8 lysates and PBS-soluble and DMSO-soluble K8 (hK8-L) were generated as described in the method section. CoCl_2_ has been widely used to induce hypoxia-mimicking conditions in the *in vitro* cell-based experiments^39,40^. In previous studies, CoCl_2_ treatment has been validated to stabilize HIF1α^39,41,42^. A bone marrow cancer cell line, SH-SY5Y or a lung cancer cell line, H460, were treated with hK8-L at the indicated concentrations or DMSO (negative control) for 12 h before treatment with 100 μM CoCl_2_ or H_2_O (negative control) for an additional 12 h incubation (**Fig. 1A**). We investigated whether these chemicals affected cell viability and proliferation at the applied concentrations. Using the water-soluble tetrazolium salt (WST) assay, the viability of SH-SY5Y and H460 cells treated with up to 100 μg/mL hK8-L, with or without 100 μM CoCl_2_, was monitored. As shown in **Fig. 1B** and **Supplementary Figure 1**, hK8-L and CoCl_2_ rarely affected cell growth and proliferation, suggesting that hK8-L was innocuous and that CoCl_2_-induced hypoxic stress rarely caused cell death at the applied concentrations.

**Fig. 1.**
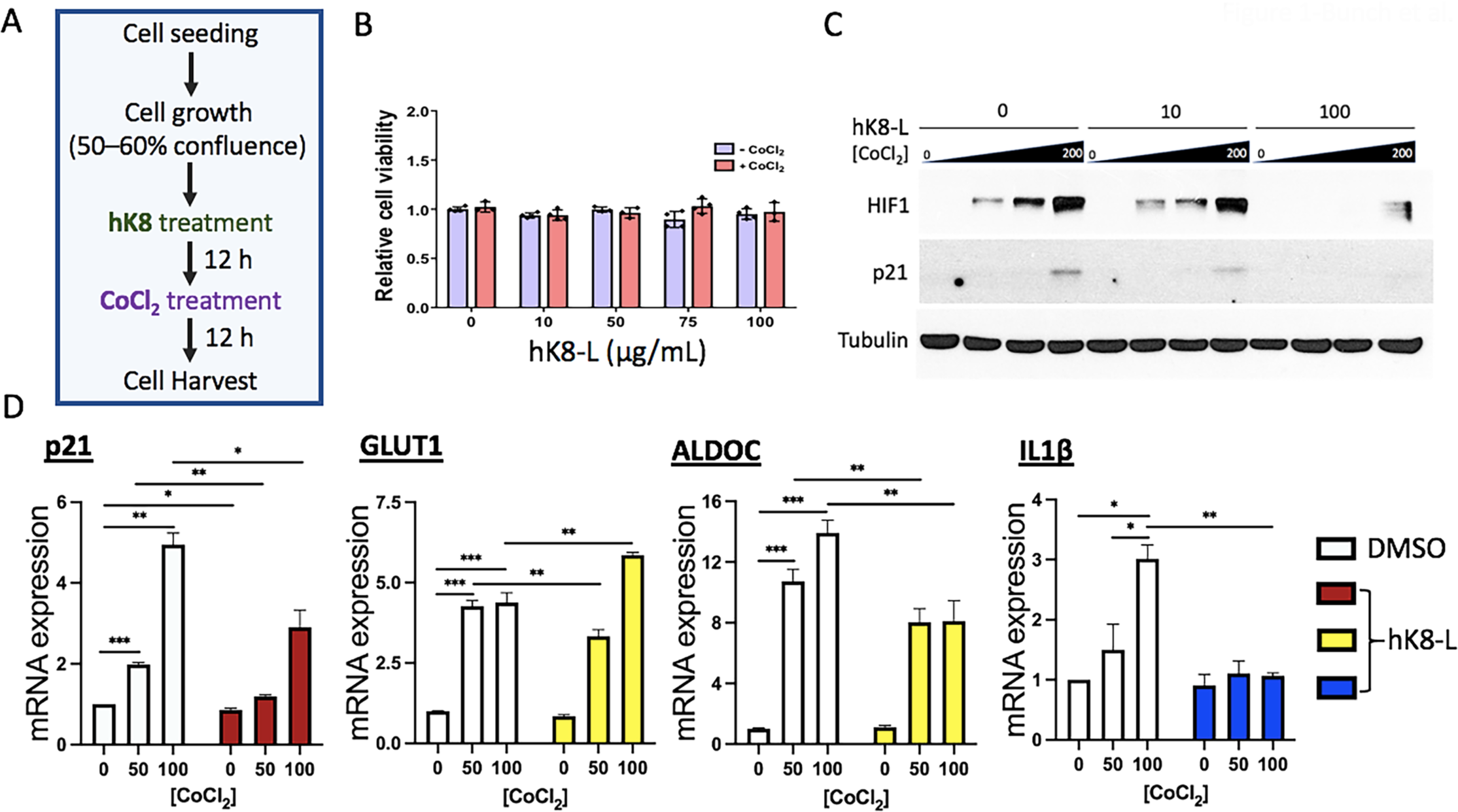
hK8-L inhibits HIF1α stabilization and its target gene expression under CoCl_2_-induced hypoxic stress. (A) A schematic representation of cell treatment with K8 lysates (0 – 100 μg/mL) and CoCl_2_ (0 – 200 μM) in this study. (B) Results of cell viability assay using the WST assay in SH-SY5Y cells. Error bars (n = 3), standard deviation (SD) throughout the figures. (C) Immunoblotting results showing the inhibitory effect of hK8-L on HIF1α (HIF1) stabilization/accumulation and p21 expression in CoCl_2_-mediated hypoxic stresses. hK8-L was supplemented to final concentrations of 0, 10, and 100 μg/mL. CoCl_2_ was treated to 0, 50, 100, and 200 μM of final concentrations. Tubulin was used as a loading control. (D) qRT-PCR results showing decreased mRNA expression of representative HIF1α-target genes, *p21*, *GLUT1*, *ALDOC*, and *IL1β* upon hypoxic stresses (0, 50, and 100 μM CoCl_2_) in the presence of 0 and 100 μg/mL hK8-L. n ≥ 3. *P < 0.05, **P < 0.01, ***P < 0.005.

Next, we investigated whether hK8-L affected hypoxic responses in SH-SY5Y cells. To answer this question, HIF1α protein levels at different doses of CoCl_2_ (0, 50, 100, and 200 μM) were quantified by immunoblotting, with 0, 10, and 100 μg/mL hK8-L (**Figs. 1A, 1C**). As expected, HIF1α protein was stabilized by CoCl_2_ treatment in a dose-dependent manner (**Fig. 1C**). Remarkably, hK8-L dramatically reduced HIF1α protein levels at 100 μg/mL and also mildly at 10 μg/mL, compared to the DMSO control. In contrast, neither the PBS-soluble K8 lysates nor 3-hydroxypropionic acid have HIF1α-suppressive effects (**Supplementary Figure 2**), suggesting that the DMSO-soluble fraction of K8 lysates is uniquely effective in alleviating HIF1α stabilization under hypoxic stresses.

### hK8-L inhibits HIF1α-target gene activation under hypoxic stresses

HIF1α is a potent transcription activator that stimulates the transcription of a number of genes in the cell and is crucial for cellular metabolic homeostasis and fate decisions. One of the HIF1α-target genes is *p21*, a cell cycle regulator^43^. Transcriptional activation at *p21* gene is well-known to be mediated by p53, an upstream transcription factor that senses genomic instability^44,45^. On the other hand, *p21* can also be activated in a p53-independent manner, and HIF1α has been reported to directly turn on the transcription of p21 by displacing MYC^43^. Therefore, the expression of *p21* with or without hK8-L and CoCl_2_ was investigated. Immunoblotting results indicated that CoCl_2_ supplementation increased p21 expression, which was positively correlated with the HIF1α levels (**Fig. 1C**). Importantly, hK8-L not only reduced HIF1α protein levels but also p21 levels in 200 μM CoCl_2_ in a dose-dependent manner (**Fig. 1C**). These results suggested that HIF1α accumulation and its transactivation at *p21* gene under hypoxic stresses are strongly inhibited by hK8-L.

Because HIF1α is a transcription factor, the transcription status of its known target genes was monitored. We selected *GLUT1* and *ALDOC*, important genes involved in glucose metabolism; *IL1β*, a critical cytokine for inflammation and immunity; and *p2*1, a potent cell cycle modulator, as these pathways are mainly upregulated by HIF1α activation^46–49^. When SH-SY5Y cells were treated with 0, 50, and 100 μM CoCl_2_ in SH-SY5Y cells, the mRNA expression of *p21*, *GLUT1*, *ALDOC*, and *IL1β* notably increased in a dose-dependent manner (**Fig. 1D**, white bars). In contrast, hK8-L supplementation at 100 μg/mL significantly interfered with the increase in all tested genes, except for *GLUT1* at the highest CoCl_2_ concentration (**Fig. 1D**, colored bars). Overall, these results suggest that hK8-L suppresses the gene expression of HIF1α target genes under hypoxic stresses.

In the consideration that K8-L might directly interfere with CoCl_2_, rather than interfering with the HIF1α stabilization, we also tested the effect of hK8-L on hypoxic responses using a different hypoxia-inducing chemical, deferoxamine methanesulfonate (DFO). hK8-L was supplemented to SH-SY5Y cells at final concentrations of 0 and 100 μg/mL and hypoxic stress was induced to the cells by the addition of DFO to final concentrations of 0, 60, and 120 μM. The protein levels of HIF1α and p21 were quantified through immunoblotting (**Fig. 2A**). The results were consistent with the CoCl_2_ data (**Fig. 1C**), showing that HIF1α was stabilized at 60 and 120 μM DFO, while hK8-L markedly suppressed the stabilization. The expression of p21 level was increased under DFO treatment, which was notably decreased in the presence of hK8-L (**Fig. 2A**). In addition, we measured the mRNA expression levels of HIF1α target genes, *GLUT1*, *ALDOC*, *p21*, and *IL1β* in the same DFO conditions, with or without hK8-L (**Fig. 2B**). The qRT-PCR data indicated the gene activation of these HIF1α target genes under DFO-induced hypoxic stresses and an overall suppression of the activation in the cells that were pre-supplemented with hK8-L, compared to the DMSO controls. This compromised HIF1α target gene expression in the presence of hK8-L in both CoCl_2_- and DFO-induced hypoxic conditions suggest that hK8-L alleviates HIF1α stabilization under cellular hypoxia, rather than interfering with CoCl_2_ or DFO.

**Fig. 2.**
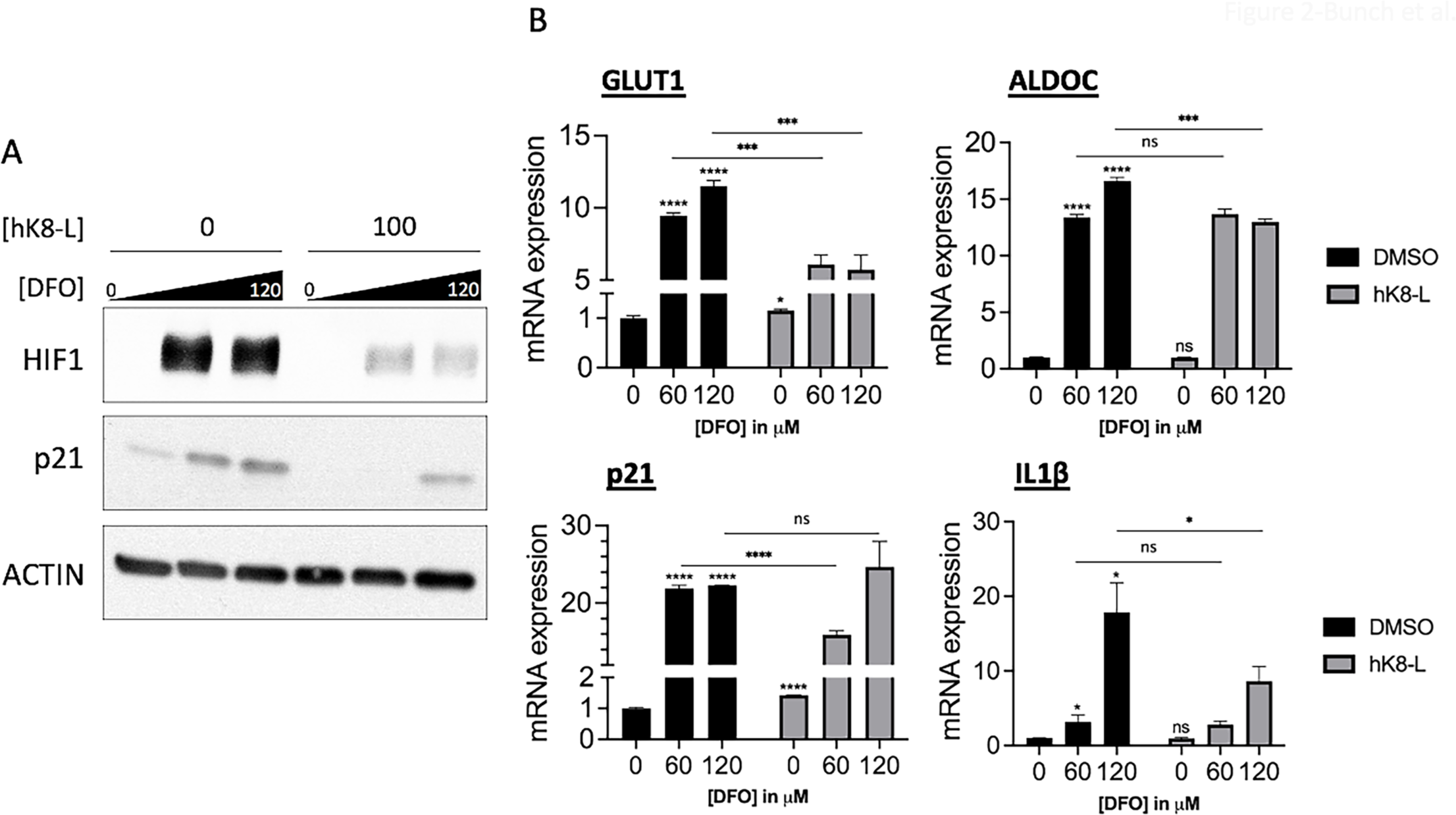
hK8-L inhibits HIF1α stabilization and its target gene expression under DFO-induced hypoxic stress. (A) Immunoblotting results showing the inhibitory effect of hK8-L on HIF1α (HIF1) stabilization/accumulation and p21 expression in SH-SY5Y cells upon DFO-induced hypoxic stress. hK8-L was supplemented to final concentrations of 0, 10, and 100 μg/mL for 12 h before DFO addition. DFO was supplemented to 0, 60, and 120 μM of final concentrations for 24 h before collecting the cells. (B) qRT-PCR results showing decreased mRNA expression of representative HIF1α-target genes, *p21*, *GLUT1*, *ALDOC*, and *IL1β* upon hypoxic stresses (0, 60, and 120 μM DFO) in the presence of 0 and 100 μg/mL hK8-L. n ≥ 3. ns, not significant, *P < 0.05, **P < 0.01, ***P < 0.005, ****P < 0.0005.

### hK8-L interferes with HIF1α and Pol II recruitment on HIF1α-target genes under hypoxia

To verify whether HIF1α destabilization is the major cause of the reduced gene expression of *p21*, *GLUT1*, *ALDOC*, and *IL1β* in the presence of hK8-L, the enrichment level of HIF1α in the transcription start sites of these genes was quantified using ChIP-qPCR. The data showed that hK8-L treatment alone either barely changed or even slightly increased HIF1α enrichment under the normoxic conditions (–CoCl_2_; **Fig. 3A**), whereas hK8-L under the hypoxic conditions (+CoCl_2_) significantly and consistently prevented HIF1α from increasing in all four genes (**Fig. 3A**). A recent study reported that hypoxia-inducible genes harbor promoter-proximally paused Pol II^50^. Therefore, Pol II occupancy was monitored in the transcription start site (TSS < ±300 nt from +1) and the gene body (GB > +301 nt from +1) of these HIF1α target genes using the target site-specific primer sets (**Table S1**) by ChIP-qPCR. Consistently with the changes in HIF1α occupancy, the results showed that the increase in Pol II occupancy in both TSS and GB under hypoxia was attenuated in the presence of hK8-L and CoCl_2_, compared to that in controls, DMSO and CoCl_2_ supplementation (**Figs. 3B, 3C**). Notably, Pol II occupancy levels were positively correlated with the levels of HIF1α enrichment levels and transcriptional activity, confirming HIF1α function as a transcriptional activator for these genes (**Figs. 3B, 3C**). These results also demonstrate the dependence of HIF1α recruitment on the transcriptional activation of *p21*, *GLUT1*, *ALDOC*, and *IL1β*, and the potent ability of hK8-L to reduce HIF1α and the transcription of these genes under the hypoxic conditions.

**Fig. 3.**
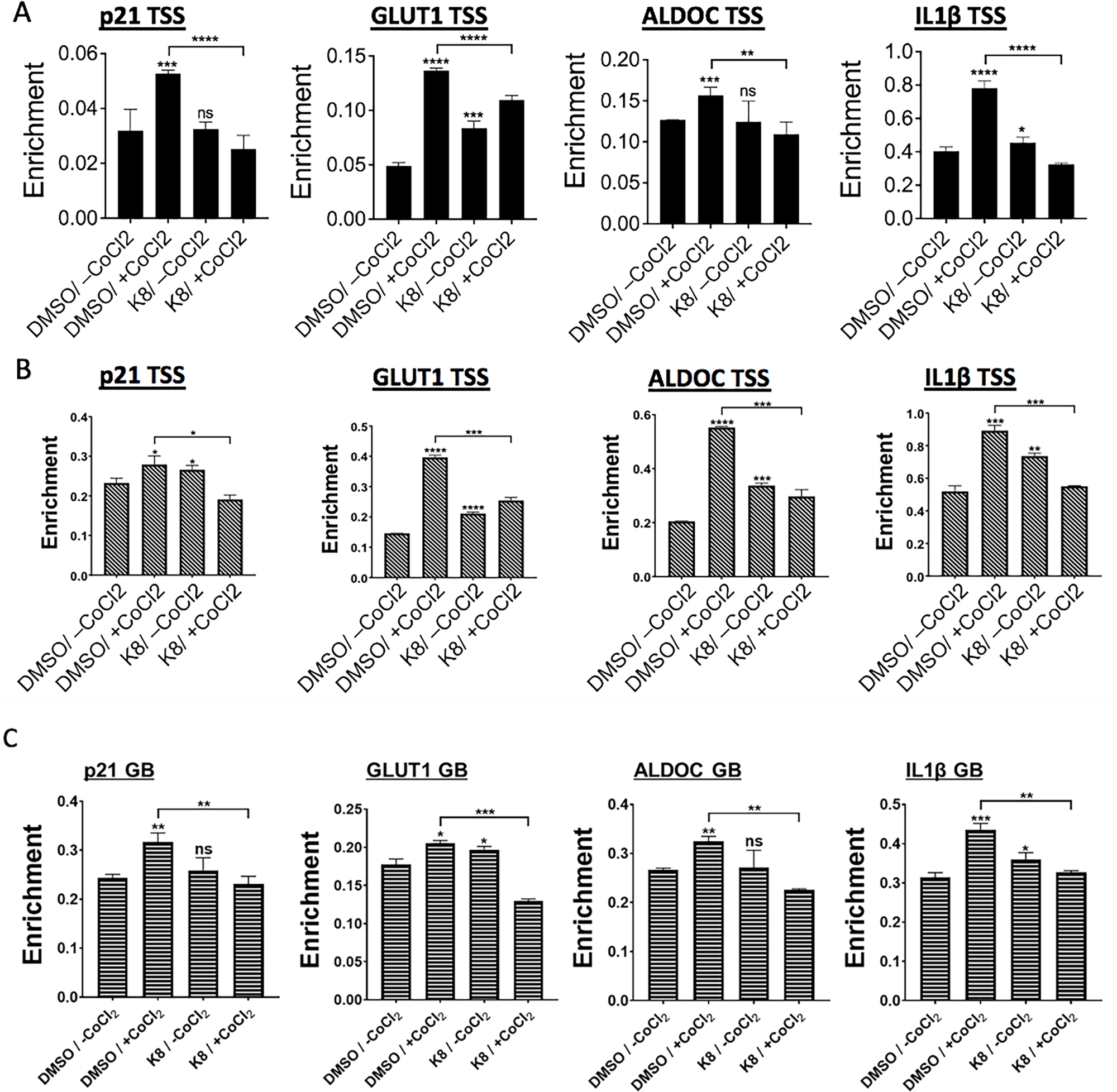
hK8-L interferes with HIF1α and Pol II recruitment under hypoxic stress. (A) ChIP-qPCR results showing HIF1α occupancy changes in the transcription start sites (TSS) of *p21*, *GLUT1*, *ALDOC*, and *IL1β* gene. Cells were treated as described in Fig. 1 before they were cross-linked and sonicated for immunoprecipitation using HIF1a antibody. n = 3. ns, non-significant throughout the figures, *P < 0.05, **P < 0.01, ***P < 0.005, ****P < 0.0005. (B) ChIP-qPCR results showing the Pol II occupancies in the transcription start sites (TSS) of *p21*, *GLUT1*, *ALDOC*, and *IL1β* gene. n = 3. *P < 0.05, **P < 0.01, ***P < 0.005, ****P < 0.0005. (C) ChIP-qPCR results showing the Pol II occupancies in the gene bodies (GB) of *p21*, *GLUT1*, *ALDOC*, and *IL1β* gene. n ≥ 3. *P < 0.05, **P < 0.01, ***P < 0.005.

### hK8-FL possesses a HIF1α-inhibitory effect similar to hK8-L

As shown **Fig. 1B**, hK8-L, the heat-inactivated K8 lysates in DMSO, excluding the PBS-soluble substances, was nontoxic to the tested cell lines. However, it is not a homogenized mixture, which includes cell debris and particles in irregular masses that are visible under a light microscope. To remove these masses and obtain a finer, completely DMSO-soluble fraction, hK8-L was centrifugated and the resulting supernatant was filtered by passing through a 0.2-μm filter. The filtered hK8-L (hK8-FL, hereafter) was tested for cytotoxicity using the WST assay. SH-SY5Y cells were treated with hK8-FL at 0, 10, 50, 75, 100, and 500 μg/mL for 12 h before supplementation with 100 μM CoCl_2_ for an additional 12 h (**Fig. 1A**). The results showed that hK8-FL was innocuous and did not affect cell viability at the tested concentrations regardless of the presence of CoCl_2_ (**Fig. 4A**). Similar results were obtained for H460 cells (**Fig. 4B**). We questioned whether hK8-FL could suppress the increase in HIF1α under hypoxic condition and if so, the inhibitory effects for the two were comparable. The level of HIF1α increase, with 200 μM CoCl_2_ treatment, was quantified by immunoblotting, comparing the cells pre-treated with DMSO, hK8-L and hK8-FL (**Fig. 4B**). The immunoblotting results indicated that both hK8-L and hK8-FL significantly reduced HIF1α protein levels in CoCl_2_-treated H460 cells and the levels of reduction with the two compounds were comparable (**Fig. 4C**). Consistently with the HIF1α results, p21 protein levels also decreased in hK8-L- and hK8-FL-treated SH-SY5Y cells, both to similar levels (**Fig. 4D**).

**Fig. 4.**
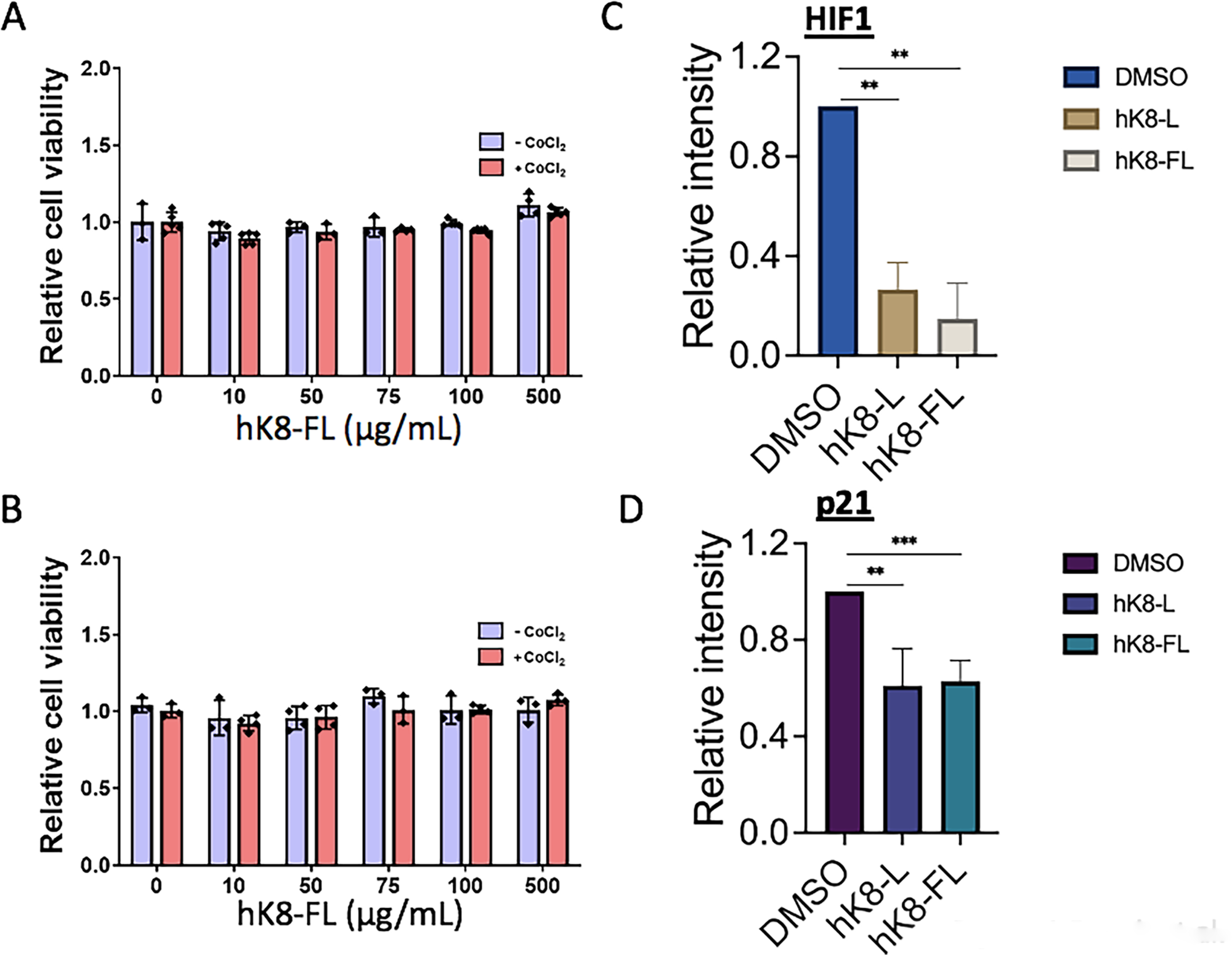
hK8-L and hK8-FL have a comparable inhibitory effect on HIF1α stabilization. (A) Results of cell viability assay using WST in SH-SY5Y cells. hK8-FL (0, 10, 50, 75, 100, and 500 μg/mL) was pre-incubated with the cells for 12 h before subjecting to the hypoxic stress induced by 100 μM CoCl_2_ (or equivalent volume of H_2_O as the control) for 12 h. n = 4. (B) Results of cell viability assay using WST in H460 cells. Same K8 and CoCl_2_ conditions were applied as **Fig. 4A**. n = 3. (C) Immunoblotting results showing the comparable inhibitory effects of hK8-L (100 μg/mL) and hK8-FL (100 μg/mL), in comparison to the DMSO control, on HIF1α accumulation during CoCl_2_ (200 μM)-mediated hypoxic stresses in H460 cells. n ≥ 3. **P < 0.01. (D) Immunoblotting results showing the comparable inhibitory effects of hK8-L (100 μg/mL) and hK8-FL (100 μg/mL) on *p21* gene activation during CoCl_2_ (200 μM)-mediated hypoxic stresses in SH-SY5Y cells. n ≥ 3. **P < 0.01, ***P < 0.005.

### Purified K8 LTA suppresses HIF1α-target gene expression

Next, K8-A, LTA purified from K8 lysates, was tested to determine whether it could suppress the hypoxic responses. Similar to hK8-L and hK8-FL, K8-A at 10, 50, and 100 μg/mL did not display cytotoxicity in SH-SY5Y cells (**Fig. 5A**). The protein level of HIF1α was measured using immunoblotting. The results indicated that unlike hK8-L and hK8-FL, the level of HIF1α after a treatment of 200 μM CoCl_2_ appeared less affected by K8-A treatment (**Fig. 5B**). However, we noticed that the HIF1α species with higher molecular weights (marked with a bracket in **Fig. 5B**) showed a decreasing trend in K8-A-treated cells, which may suggest certain changes in the post-translational modifications or the dimerization status of HIF1α by K8-A. The mRNA expression of *p21*, *ALDOC*, and *GLUT1* significantly decreased under hypoxic stresses in K8-A-treated cells, as well as under normoxic conditions, except for *p21* at 50 μg/mL K8-A (**Figs. 5C–E**).

**Fig. 5.**
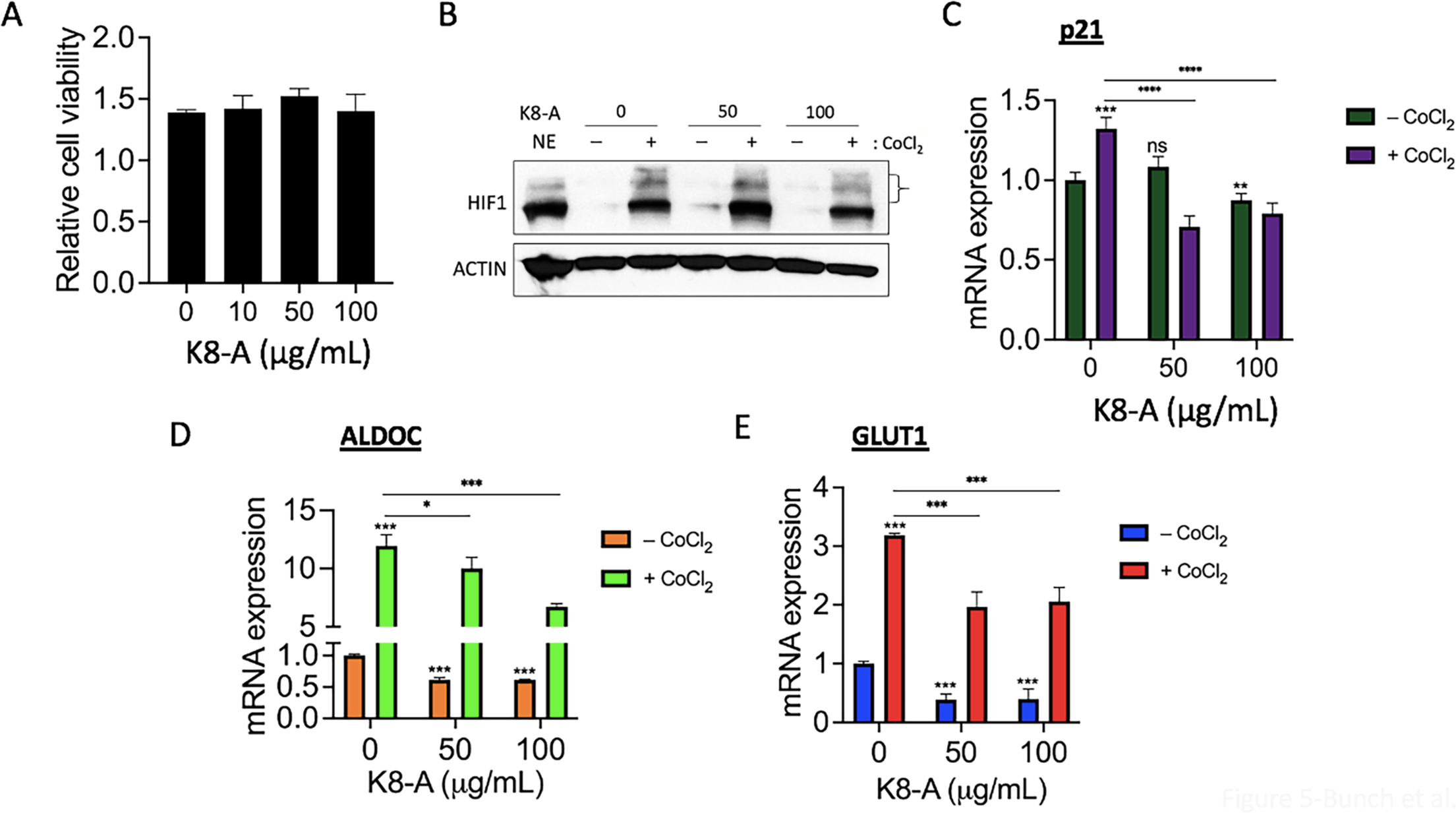
K8-A inhibits HIF1α-target gene activation. (A) Results of cell viability assay using WST in SH-SY5Y cells. K8-A was supplemented to final concentrations 0, 10, 50, and 100 μg/mL and the cells were incubated for 24 h. n = 3. (B) Immunoblotting results showing the effect of K8-A on HIF1α protein in SH-SY5Y cells. K8-A was treated to final concentrations of 0, 50, and 100 μg/mL. CoCl2 was treated to a final concentration of 200 μM. Same time points as Fig. 1A were applied for K8-A experiments. β-ACTIN was used as a loading control. NE, HeLa NE as a technical reference. HIF1α species with higher molecular weights marked with a bracket. (C) qRT-PCR results showing the effect of K8-A on the mRNA expression of *p21*, a HIF1α-target gene during normoxic and CoCl_2_ (200 μM)-mediated hypoxic conditions in SH-SY5Y cells. n ≥ 3. **P < 0.01, ***P < 0.005, **** < 0.0005. (D) qRT-PCR results showing the effect of K8-A on the mRNA expression of *ALDOC*, a HIF1α-target gene during normoxic and CoCl_2_ (200 μM)-mediated hypoxic conditions in SH-SY5Y cells. n ≥ 3. *P < 0.05, ***P < 0.005. (E) qRT-PCR results showing the effect of K8-A on the mRNA expression of *GLUT1*, a HIF1α-target gene during normoxic and CoCl_2_ (200 μM)-mediated hypoxic conditions in SH-SY5Y cells. n ≥ 3. ***P < 0.005.

### hK8-L inhibits the gene expression of HIF1α regulators, PHD2 and VHL

Lastly, we analyzed the mRNA expression levels of prolyl hydroxylase domain protein 2 (*PHD2*, also known as EGLN1) and *VHL*, two proteins that regulate HIF1α stability and hypoxic responses, using qRT-PCR. In the control (DMSO) group, mRNA levels of *PHD2* and *VHL* genes were increased and decreased upon 200 μM CoCl_2_ treatment in SH-SY5Y cells, respectively (**Fig. 6A**). The cells were supplemented with 100 μg/mL hK8-L or hK8-FL prior to CoCl2 treatment. The expression of *PHD2* and *VHL* mRNAs decreased, compared to that in the control (**Fig. 6A**). *PHD2*, both a regulator and target gene of HIF1α^15,16^. The PHD2 protein was quantified using immunoblotting. Consistently with the qRT-PCR results, hK8-FL treatment decreased the PHD2 protein levels (**Fig. 6B**). Overall, these data suggested that DMSO-soluble K8 lysates have the effect to inhibit HIF1α activity in SH-SY5Y cells.

**Fig. 6.**
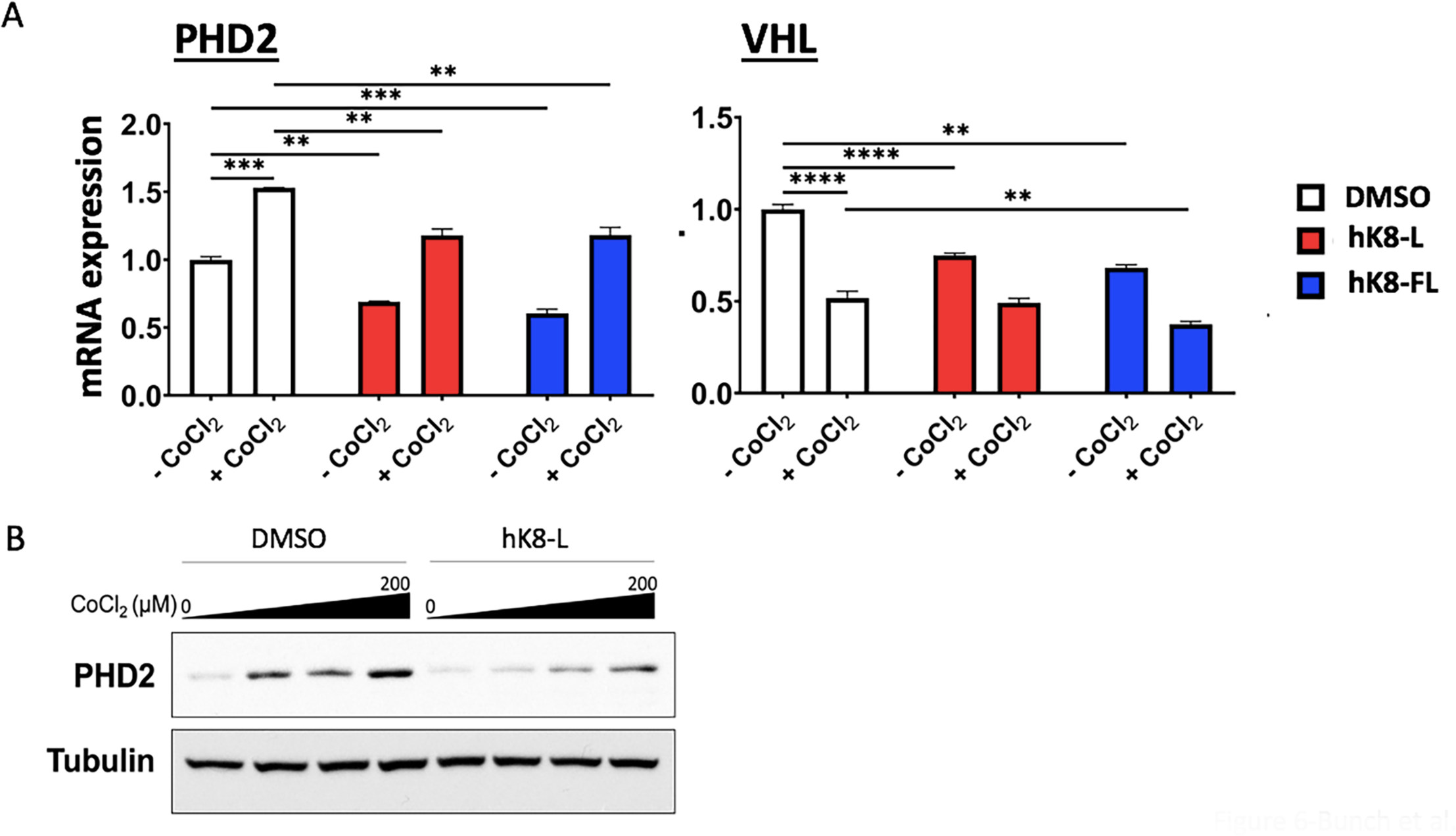
hK8-L inhibits HIF1α-regulators, *PHD2* and *VHL*. (A) qRT-PCR results showing the effect of hK8-L (100 μg/mL) and hK8-FL (100 μg/mL) on the mRNA expression of *PHD2*, a HIF1α-regulator and -target gene and *VHL* during normoxic (-CoCl2) and CoCl_2_ (200 μM, + CoCl_2_)-mediated hypoxic conditions in SH-SY5Y cells. n ≥ 3. **P < 0.01, ***P < 0.005, ****P < 0.0005. (B) Immunoblotting results showing the effect of hK8-L on *PHD2* protein levels in SH-SY5Y cells. hK8-L was treated to final concentrations of 0 (DMSO only) and 100 μg/mL. CoCl2 was treated to final concentrations of 0, 50, 100, and 200 μM. α-Tubulin was used as a loading control.

## DISCUSSION

It is important to note that hypoxic responses occur in inflammation and cancer growth. Chronic inflammation often develops into cancers. For example, people with chronic inflammatory bowel diseases such as Crohn disease are exposed to a higher risk to colon cancers^51,52^. It is well-known that HIF1α is stabilized also in normoxia during inflammation^53^. While chronic inflammation shows the increased level of HIF1α, a study reported that HIF1α is required for the resolution of bowel inflammation^54^. Another study reported that HIF1α aggravates psoriasis by inhibiting BMP6 expression^55^. As the physiological roles of HIF1α for hypoxic responses, inflammation, and immunity are important, the molecules to control the HIF1α level have been developing and clinically tried^56^. The identification of the novel function of lactobacillus K8 extracts to modulate HIF1α stability in this study suggests a way to develop a potential natural remedy to treat HIF1α deregulation, which is implicated in different disease pathologies.

In this study, we tested *Lactiplantibacillus plantarum K8* lysates on the human hypoxic responses *in vitro*. This property of K8 lysates is pertained to the DMSO-soluble fractions of the freeze-powdered K8 lysates (hK8-L and hK8-FL). Both hK8-L and hK8-FL significantly inhibit HIF1α activation, thereby interfering with the HIF1α-mediated hypoxic gene expression (**Fig. 1–4**). Representative HIF1α-target genes, including *p21*, *ALDOC*, *GLUT1*, and *IL1β,* were shown to be downregulated under hypoxic stresses in the presence of hK8-L (**Figs. 1–3**). The word “modulation” seems appropriate to describe the K8 results because hK8-L slightly increases the mRNA synthesis and HIF1α/Pol II occupancy of some genes in normoxia, while it inhibits the increase in hypoxia (**Figs. 1D, 3A–C**). These data suggest that hK8-L may slightly stimulate hypoxic responses including glucose uptake (GLUT1) and metabolism (ALDOC) under normoxic conditions, while it functions as a break to prevent these genes from being acutely transcribed in hypoxia.

In addition, hK8-FL and K8-A exhibited an inhibitory effect on hypoxia-induced gene expression (**Figs. 4, 5**). Filter cleared fraction of hK8-L (hK8-FL) is void of DMSO-insoluble substances larger than 0.2 μm. Under microscopic examination, we noted that centrifugally cleared hK8-L included a number of irregular sized, visible substances, thus not homogenized. Thus, the homogenzed hK8-FL data showing the suppressive effect on HIF1α is meaningful in that it confirms the effects to be contributable to the DMSO-soluble, small compounds in K8 cells. In addition, we qualitatively and quantitatively analyzed hK8-L and PBS-K8 using gas chromatography and mass spectrometry (GC/MS)(**Table S2–4**; **Supplementary Data 1–3**). Overall, in the analysis of volatile components, K8 powder (52 compounds), hK8-L (24 compounds), and PBS-K8 (33 compounds) don’t share compounds with one another. Between hK8-L and PBS-K8, only two compounds were commonly identified. Because there are no chemical libraries for K8 to validate and compare with our GC/MS data, it is difficult to pinpoint certain chemicals that might contribute to the suppressive effect on HIF1α stabilization among the compounds. However, our GC/MS data verified chemically-differentiated K8 fractions by two solvents, PBS and DMSO and DMSO to efficiently extract the effect molecule(s) from K8 powder.

Initially, we hypothesized that VHL may be increased by hK8-L pretreatment before hypoxic stress. This hypothesis was based on the fact that VHL is an E3 ligase for HIF1α for degradation^5,13,18^, which may explain how HIF1α is decreased in the presence of hK8-L in spite of hypoxic stresses. Instead, qRT-PCR results showed a decrease in *VHL* expression in the presence of hK8-L and hypoxic stress (**Fig. 5A**). This suggests that the K8-mediated HIF1α protein reduction is probably not through the PHD–VHL degradation pathway but through other mechanisms. Further studies are required to identify the exact underlying mechanisms: how K8 lysates inhibit HIF1α protein accumulation under hypoxic stresses. In addition, transcriptome analysis is necessary for identifying all hypoxic genes regulated by hK8-L/FL and for understanding the impact of these compounds on hypoxic and overall gene regulation.

Together, our data support our hypothesis that K8 cellular compounds, hK8-L/FL and K8-A regulate hypoxic responses though controlling HIF1α in SH-SY5Y and H460 cells. This study suggests that hK8-L/FL and K8-A inhibit the accumulation of HIF1α and repress the expression of representative HIF1α target genes under hypoxic stresses. Our data provide the first pieces of evidence that natural compounds in lactobacillus K8 lysates have the potential to be developed into a hypoxic regulator. We propose that hK8-L/FL may provide health benefits to humans as a natural modulator/suppressor of hypoxic responses, which are critical for cancer cell proliferation and inflammation.

## Acknowledgments

We thank C. Jung, M. Seu, and the current members of the Bunch laboratory members at Kyungpook National University (KNU) for their technical assistance. H.B. thanks J. Christ and John for their loving encouragement and support throughout the course of this work.

## Financial Supports

This research was supported by grants from the National Research Foundation (NRF) of the Republic of Korea (2022R1A21003569) to H.B.

## Author Contributions

SJ, ND, JL, JJ, DK, CR, and HB performed qRT-PCR. BK, JJ, and HB carried out immunoblotting. JJ, DK, and HB performed ChIP-qPCR. HB and JJ generated PBS-K8, hK8-L and hK8-FL. NP and D(-H)C carried out FACS. HK and D(K)C provided freeze-dried K8 powder and research materials. HB created the hypothesis, designed the experiments, analyzed and curated the data, and wrote and revised the manuscript.

## Data and Materials Availability

All data are available in the manuscript or as supplementary information.

